# A distinctive subset of microglia positioned at the paraventricular zone are dedicated to cleaning the cerebrospinal fluid

**DOI:** 10.1101/2025.02.23.639717

**Authors:** Guoyang Zhou, Jian He, Baojie Mao, Guo Cheng, Rui Zhang, Qiang Wang, Xiaoli Liu, Jiaxin Zheng, Ming Wang, Li Zhao, Peng Shi, Xiao Z Shen, Shu Wan

**Affiliations:** Brain Center, Affiliated Zhejiang Hospital, Zhejiang University School of Medicine, Hangzhou, Zhejiang, China; Zhejiang Key Laboratory of Geriatrics and Geriatrics Institute of Zhejiang Province, Zhejiang Hospital, Hangzhou, Zhejiang, China; Research Center for Life Science and Human Health, Binjiang Institute of Zhejiang University; Department of Cardiology, The Second Affiliated Hospital, School of Medicine, Zhejiang University, Zhejiang, China; Department of Physiology and Department of Cardiology of the Second Affiliated Hospital, Zhejiang University School of Medicine, Hangzhou, Zhejiang, China; Department of Laboratory Medicine, Affiliated Zhejiang Hospital, Zhejiang University School of Medicine, Hangzhou, Zhejiang, China; Department of Neurology, Affiliated Zhejiang Hospital, Zhejiang University School of Medicine, Hangzhou, Zhejiang, China; Key Laboratory for Biomedical Engineering of Ministrey of Education, Collage of Biomedical Engineering & Instrument Science, Zhejiang University, Hangzhou, Zhejiang, China; Institute of Translational Medicine, Zhejiang University School of Medicine, Hangzhou, Zhejiang, China; State Key Laboratory of Transvascular Implantation Devices, Hangzhou, Zhejiang, China

## Abstract

The interconnected ventricles in the central nervous system are filled with cerebrospinal fluid (CSF) which serves multifaceted roles including bring away metabolic wastes. However, whether there is an inherent “dislodging” mechanism, in addition to CSF drainage and reabsorption, at play for the removal of microparticles in the CSF is unclear. In this study, we identified a subset of microglia distributed in close proximity to the ependymal wall of ventricles. They differed from parenchymal microglia in morphology, behaviors and transcriptomes. In particular, the paraventricular microglia extended transependymal dendrites into the lumen of ventricles and were proficient in sequestering and phagocytosing intraventricular exogenous particles when present. A specific removal of the paraventricular microglia led to an acute ventriculomegaly due to an increased colloid osmotic pressure in the CSF resulting from protein accumulation. Thus, we identified the paraventricular microglia as specific guardians in monitoring and cleaning the CSF.

## INTRODUCTION

The brain contains an interconnected system of ventricles filled with cerebrospinal fluid (CSF). The ventricle wall comprises a specific epithelium sheet of ciliated ependymal cells. Unlike most epithelial cells, ependymal cells are not underlined by a basal lamina and do not form tight junctions between each other ^1^, which renders a relatively easy exchange between interstitial fluid and CSF. Choroid plexuses are considered the main site of CSF production, with extra-choroidal portion mainly originated from the interstitial fluid^2^. The CSF flows in a direction from the lateral to the third and fourth ventricles, and the major outlet of CSF is the subarachnoid space where it is resorbed by arachnoid villi and granulations. CSF serves multifaceted roles in removing metabolic wastes from the brain, maintenance of environmental physicochemical homeostasis, and regulation of intracranial pressure balance, etc ^2^. CSF consists predominantly of water carrying various solutes including electrolytes, proteins, neurotransmitters ^3^. The volume of CSF is at least partly dependent upon hydrostatic and osmotic gradients ^4^. CSF can change composition as a function of age or pathophysiological states including abnormal brain activities. For example, lactate concentration was elevated following motor-onset epileptic seizures ^5^; amyloid and tau proteins could rise even before symptom onset of Alzheimer’s disease ^6,7^. One of the mechanisms underlying ageing and AD is thought to be a reduced interstitial fluid-CSF exchange, leading to the pathogenic accumulation of amyloid and tau ^8,9^. Thus, a consistent CSF flow and the associated clearance is important for brain health. However, whether there is an inherent cellular “dislodging” mechanism, in addition to CSF drainage and reabsorption, at play for the removal of pathological particles when incremented in the CSF is unclear.

Resident macrophages are the most populated immune cells in the central nervous system (CNS), consisting of microglia residing in the parenchyma and macrophages in the borders including perivascular space, choroid plexus, and meninges ^10^. Given their specific niches, these phagocytes are independent from each other in adults and assume specific functional interaction with their surrounding cells and proximal anatomic structures ^11^. In this study, we identified a subset of microglia populated in close apposition to the ventricle ependymal wall. They differ in morphology, behavior and transcriptome from their parenchymal microglial counterparts, and specialized in cleaning sedimentary particles in the CSF. Without them, acute ventriculomegaly occurred as a result of an increased colloid osmotic pressure in the CSF due to protein accumulation.

## RESULTS

### There is a population of paraventricular phagocytes distributing on the walls of brain ventricular system

*Cx3cr1*^GFP/+^ reporter mice could label both parenchymal microglia and border macrophages ^12^. We noticed that in the areas within 50 μm from the lateral ventricular wall were underrepresented with GFP^+^ phagocytes relative to the nearby brain parenchyma (Figure 1A). The underrepresentation of phagocytes around the lateral ventricular wall was further confirmed by Iba1 staining in adult C57BL/6 mice (Figure S1A). Distinguishably, most of the phagocytes in this paraventricular zone (within 50 μm) were in close proximity to the AQP4^+^ ependymal wall of the ventricle (Figure 1A) (The remaining AQP4^+^ components in the parenchyma were astrocyte endfeet surrounding capillaries). This distribution pattern of paraventricular phagocytes was also evident in the 3^rd^ and 4^th^ ventricles (Figure S1B). Moreover, these paraventricular phagocytes displayed a distinctive morphology from the parenchymal microglia, as they had amoeboid morphologies with short and stunted processes, in contrast with parenchymal microglia that had ramified shapes with long, thin processes (Figures 1A and 1B)^13^. Moreover, the number of processes per cell was markedly lower in these phagocytes (Figure 1B) with a decreased process complexity as manifested by Sholl analysis (Figure 1C). Confocal analysis of thicker sections (30 μm) of the ventricular wall area with high-resolution and optical magnification revealed that the paraventricular phagocytes engaged in tight contacts with AQP4^+^ ependymal cells, as all of them formed transependymal processes (Figure 1A, ①). Besides, some phagocytes actually embedded cell bodies in the ependymal wall (Figure 1A, ②) and some cells have migrated to the apical side of the ependymal wall (Figure 1A, ③). Three-dimensional (3D) reconstruction revealed that many transependymal processes derived from the paraventricular phagocytes were actually exposed to the ventricular lumen (Figure 1D), which was further confirmed by co-staining with β-catenin, a marker of adherent junction associated with ependymal layer (Figure 1E).

**Figure 1.**
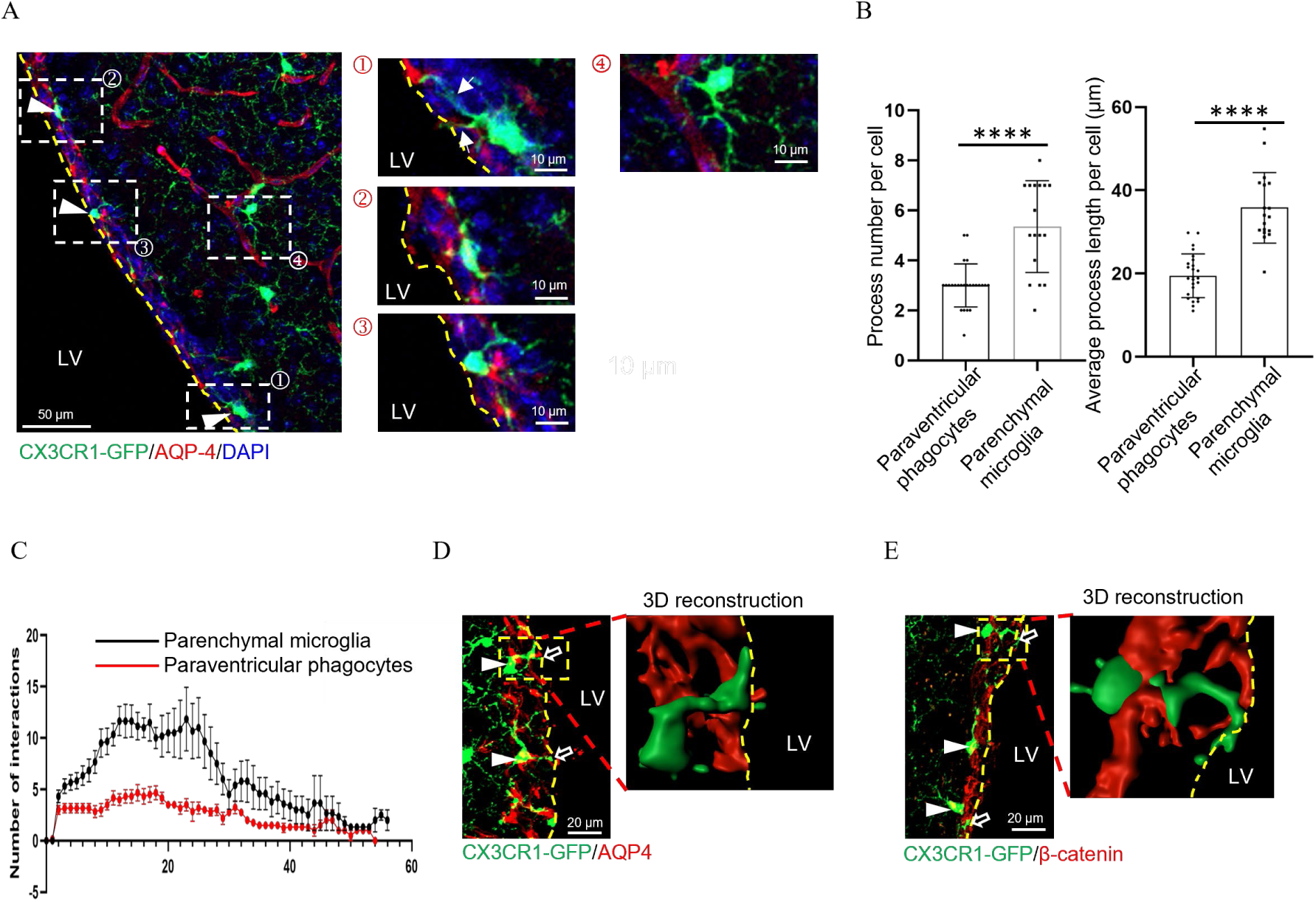
The morphology of paraventricular phagocytes. (**A**) CX3CR1^+^ cells are distributed in the paraventricular zone of reporter *Cx3cr1*^GFP/+^ mice. Co-staining for AQP4 is shown. Representative images from 5 mice of each group. Arrowhead, paraventricular phagocyte. ①, a paraventricular phagocyte formed a transependymal process (arrow); ②, a paraventricular phagocyte was embedded in the ependymal wall; ③, a paraventricular phagocyte is present in the apical side of the ependymal wall; ④, a typical microglia in brain parenchyma. Maximum z-projection of 25 μm. (**B**) Quantification of protrusion number and mean protrusion length per paraventricular phagocyte or parenchymal microglia. Each dot indicates the quantification of one cell. Data were pooled from 6 mice. (**C**) Sholl analysis of branch ramification complexity of all paraventricular phagocytes and parenchymal microglia used in (B). (**D** and **E**) Confocal image of the spatial relationship between AQP4^+^ ependymal cells (D) or β-catenin^+^ ependymal wall and CX3CR1^+^ cells (GFP). The magnifications show the corresponding 3D reconstructions. (D and E) Arrowhead, paraventricular phagocyte; arrow, the process derived from the paraventricular phagocytes were exposed to the ventricle lumen. The apical side of ventricular wall is indicated with dashed line. LV, lateral ventricle. \*\*\*\**P*<0.001 by two-tailed unpaired t test. Data are depicted as mean±SEM. Data are derived from at least 2 independent experiments.

### Interactions between the paraventricular phagocytes and the ventricular ependymal wall

To investigate the dynamics of interactions between paraventricular phagocytes and the ependymal wall, live brain slices were made from *Cx3cr1*^GFP/+^ mice. The slices were 150 μm in thickness, much larger than the diameter of soma of either phagocytes (∼8 μm) or ependymal cells (∼5 μm). We tracked the cells by two-photon microscope for up to 2.5 h in the depth of 75 μm, a place minimally disrupted by slice preparation. Interestingly, while the processes of parenchymal microglia moved in all directions with a low speed (Figure 2A and Video S1), there was a polarity of paraventricular phagocytes in process movement, with the processes protruding towards the ventricles make movement with a higher frequency, appearing sampling the ventricle contents and probing intraventricular environment (Figure 2A, and Videos S2-S3). In addition, while the soma of parenchymal microglia remained sessile and fixed in position during the imaging session, some paraventricular phagocytes were observed to make transmigration either from the basolateral side of the wall to the apical side or vice versa (Figures 2B and 2C, and Videos S4 and S5). Moreover, we observed crawling phagocytes that lost polarity at the apical CSF-containing surface of the ventricle (Figure 2D and Video S6), reminiscent of supraependymal cells ^14^. When fluorescent latex beads were injected into the ventricular space, a dynamic association and take-up of inert beads by the paraventricular phagocytes occurred (Figure 2E), while neither parenchymal microglia (Figure 2E, left) nor choroid plexus stromal macrophages (CpMØ) (Figure 2E, right) had close contact with the intracerebroventricularly (ICV) infused beads at any given time. A similar phagocytosis of red blood cells (RBCs) by the paraventricular phagocytes was also observed when RBCs were ICV injected (Figure 2F). Thus, paraventricular phagocytes were dynamically interplaying with the ventricular ependymal wall and intaking intraventricular particles.

**Figure 2.**
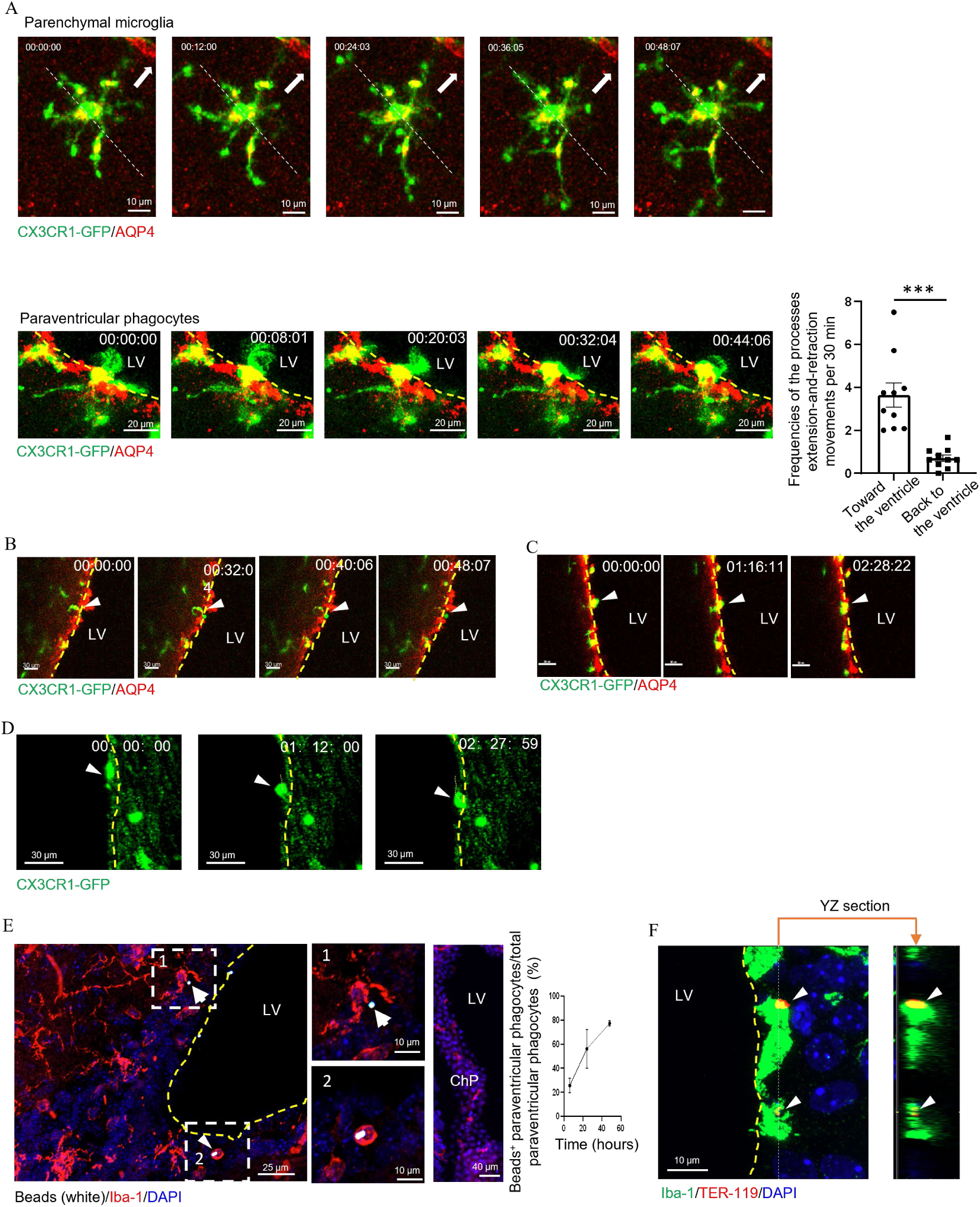
Paraventricular phagocytes dynamically move their transependymal processes and “sample” intraventricular particles. (**A**) Time lapses of parenchymal microglia (upper) and paraventricular phagocytes (lower). Frequencies of the processes facing toward and back to the ventricle in making extension-and-retraction movements are quantitated. Each dot indicates one cell and a total of 3 mice were examined. Arrow in the upper panels indicates the direction towards ventricles. (**B** and **C**) Time lapses of representative paraventricular phagocytes (GFP) moving from the basolateral side to the apical side of the AQP4^+^ ependymal wall (B) or vice versa (C). (**D**) Time lapses of a paraventricular phagocyte was crawling along the apical side of the ventricle wall. (A-D) Z-projections of 30 μm. (**E**) A representative picture shows at 6 hr post ICV injection of fluorescent beads, some beads (white) had been phagocytosed by paraventricular phagocytes (red). A kinetics panel shows that bead-associated paraventricular phagocytes increased in number along the observation period. ChP, choroid plexus. (**F**) At 48 hr post ICV injection of red blood cells (red), their space relationship with paraventricular phagocytes (green) is shown. Dashed line, the border of ventricle wall. LV, lateral ventricle. ns. not significant. \*\*\*\**P*<0.001 by two-tailed unpaired t test. Data are depicted as mean±SEM. Data are derived from at least 2 independent experiments.

### The paraventricular phagocytes are a subpopulation of microglia

The CNS is distributed with parenchymal microglia and border macrophages. To probe the identity of these paraventricular phagocytes, we first examined a plethora of markers widely used to differentiate microglia from CNS border macrophages. The paraventricular phagocytes expressed microglia markers P2Y12 and TMEM119 ^15^ which are not expressed by border macrophages including CpMØ (Figures 3A and 3B). Instead, the paraventricular phagocytes, as well as parenchymal microglia, did not express CD206 which was present on nearby CpMØ and perivascular macrophages (Figures 3C and S2A) ^16^. LYVE-1 was reported to be expressed by perivascular and meningeal macrophages but not microglia or CpMØ ^17,18^. We could discern this expression pattern of LYVE-1 in the CNS (Figures 1D and S2B). The paraventricular phagocytes did not express LYVE-1 (Figure 3D). Altogether, homeostatic surface markers point out that the paraventricular phagocytes are more akin to microglia over border macrophages.

**Figure 3.**
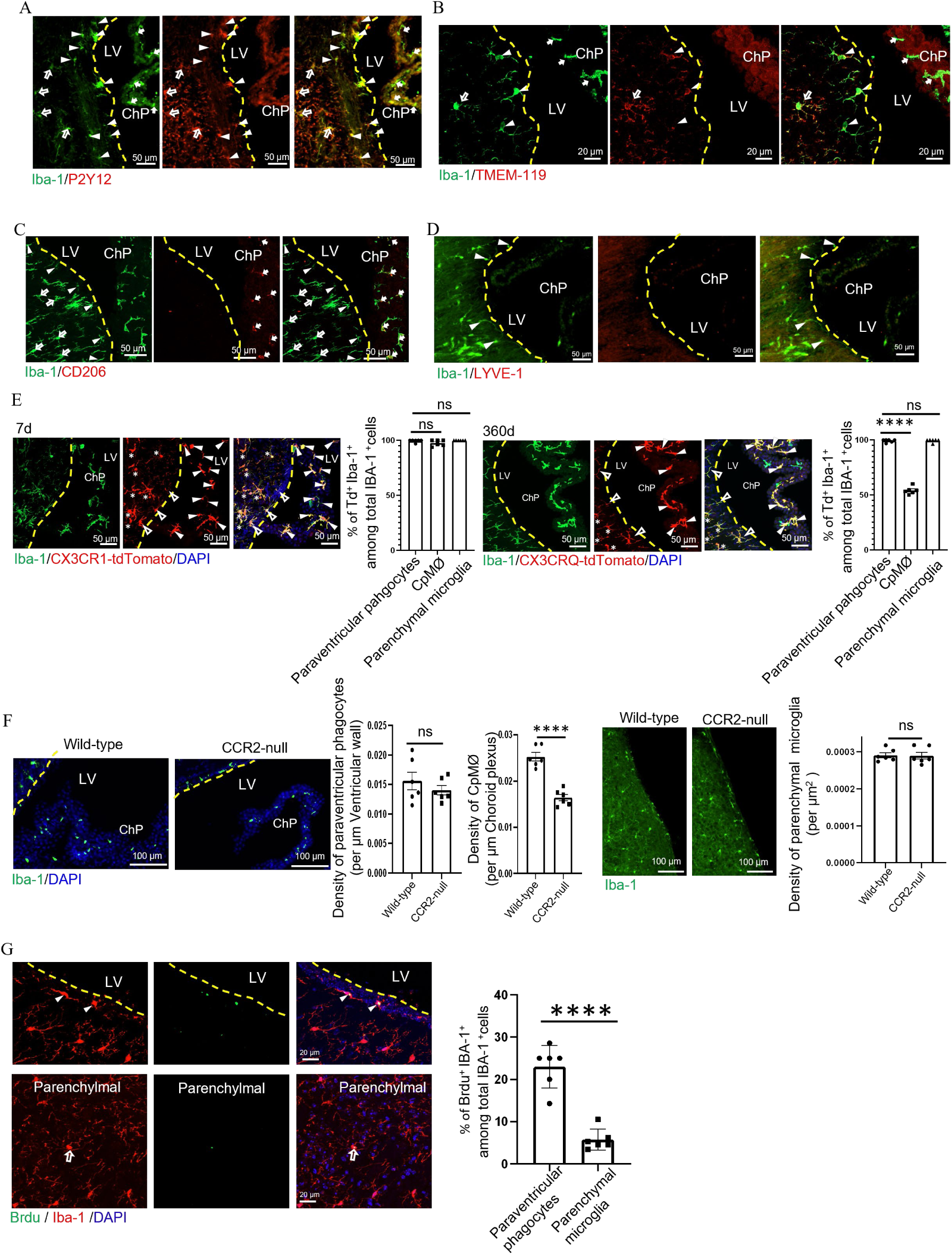
The paraventricular phagocytes are a specific subset of microglia. (**A**-**D**) Representative pictures show P2Y12 (A), TMEM119 (B), CD206 (C) and LYVE-1 (D) expression in paraventricular phagocytes (arrowheads), parenchymal microglia (hollow arrows) and CpMØ (solid arrows). (**E**) 8-weeks old *Cx3cr1*^CreERT2^: *Rosa26*^tdTomato^ mice were i.p. treated with tamoxifen. The percentages of tdTomato^+^ cells in paraventricular phagocytes (hollow arrowheads), parenchymal microglia (asterisks) and CpMØ (solid arrowheads) were traced on day 7 and day 360. (**F**) 3-month-old CCR2-null and wild-type littermates were examined for the densities of the indicated phagocytes. (**G**) The frequencies of BrdU^+^ cells among paraventricular phagocytes (arrowheads) and parenchymal microglia (arrows). Dashed line, the border of ventricle wall. ChP, choroid plexus. ns. not significant. \*\*\*\**P*<0.001 by two-tailed unpaired t test (F and G) and one-way ANOVA test (E). Data are depicted as mean±SEM. Data are derived from at least 2 independent experiments.

Next, we adopt a fate mapping strategy via employing *Cx3cr1*^CreERT2^: *Rosa26*^tdTomato^ mice. Tamoxifen treatment induced a near full labeling of parenchymal microglia, paraventricular phagocytes and the nearby CpMØ (Figure 3E). Mice were then tracked for one year after labeling. Previous fate mapping studies revealed that microglia are maintained by self-renewal, while constant monocyte trafficking to the choroid plexus near ventricles throughout adult life to replenish resident CpMØ ^12^. Concordant with their maintaining patterns, quantitative examination on histological slices showed constant high expression of the reporter gene in microglia; in contrast, tdTomato-labeling progressively decreased in Iba-1^+^ CpMØ, suggesting a dilution by blood-borne cells (Figure 3E). The labelling rate of paraventricular phagocytes did not drop and was comparable to parenchymal microglia over the 360-d observation window (Figure 3E). Substantiating the notion that the paraventricular phagocyte is a subpopulation of microglia but not relies on monocytes replacement, *Ccr2*-deficient adult mice had normal distribution of parenchymal microglia and paraventricular phagocytes, while CpMØ were reduced (Figure 3F)^12^. The notion of their self-renewal was further strengthened by a 22.98% of proliferating rate when examined by BrdU incorporation (Figure 3G). The proliferating rate of paraventricular phagocytes was even higher than parenchymal microglia (∼5.76%) (Figure 3G), suggesting a dynamic replenishment for their likely loss by transmigating into the ventricles, which was consistent with the previous observations of the presence of microglia in normal CSF^19^.

Taken together, the characteristics of signature markers, fate mapping and local proliferation suggest that the paraventricular phagocytes were a subpopulation of microglia distributing in a specific niche, i.e., the border of ventricles. We designated them as paraventricular microglia (PvtMG) hereafter.

### Paraventricular microglia exhibit a distinct transcriptome signature

To further investigate the roles of the ependyma-associated PvtMG, we next performed transcriptome analysis. To this end, 200-μm-thick brain slices including lateral ventricle derived from *Cx3cr1*^GFP/+^ mice were prepared. The ependymal barrier had high stiffness relative to the surrounding brain parenchyma due to its enrichment with collagen (Figure S3A). Thus, a ∼30-50 μm-width slice of tissue including ependymal wall could be easily separated by using a 33-gauge needle. After a brief mechanical dissociation within artificial cerebrospinal fluid, GFP^+^ ependyma-associated PvtMG were sorted followed by bulk RNA-sequencing (RNA-seq) (see Methods). To evaluate the characteristics of PvtMG in a strict manner, similar numbers of microglia derived from basal ganglion (BG, a brain region close to ventricle) and cerebrocortex (CTX, a brain region far from ventricle) of the same batches of mice were used for transcriptomics comparison (Figure 4A).

**Figure 4.**
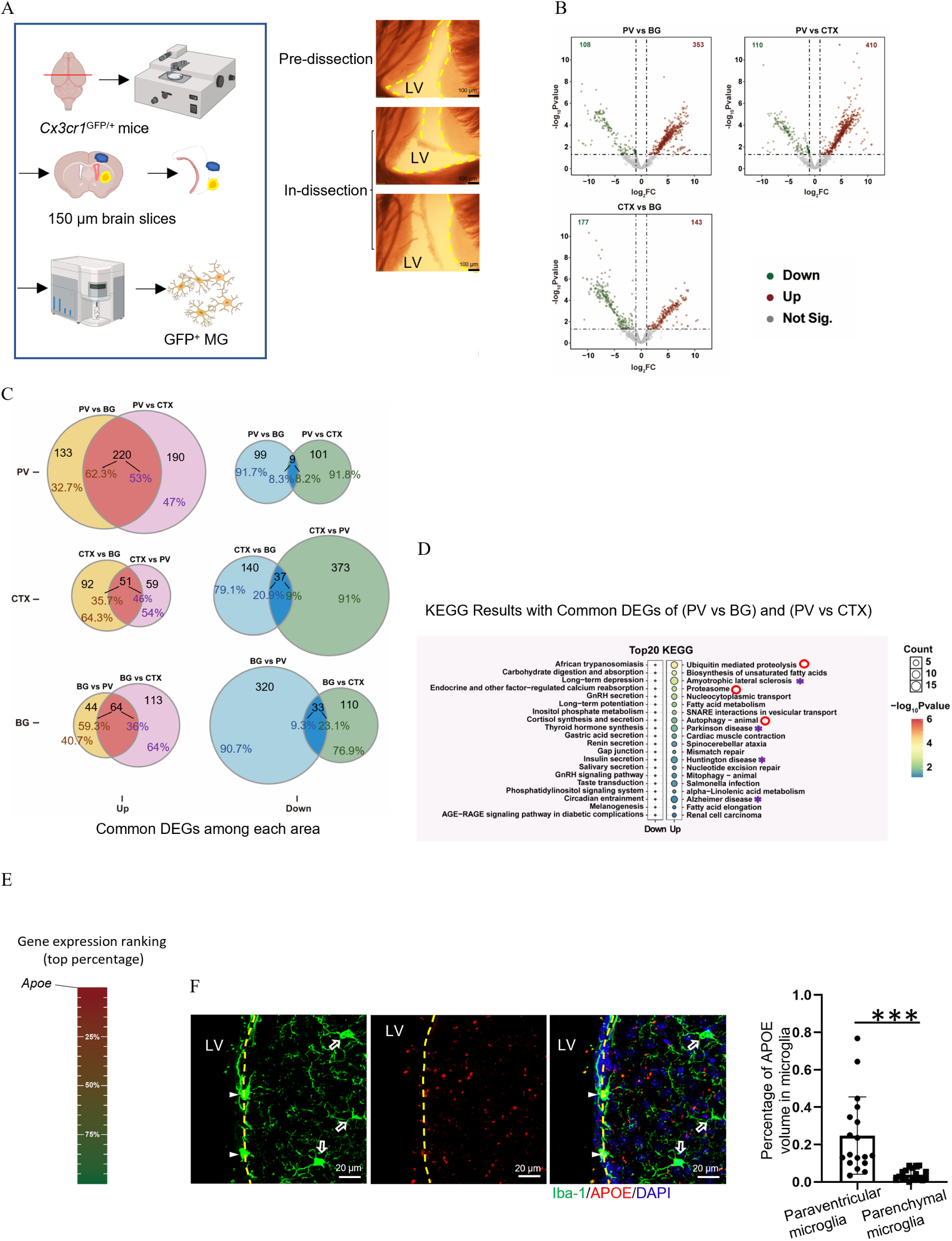
Transcriptomic difference between PvtMG and parenchymal microglia. (**A**) Schematic outline for transcriptome analysis of dissociated microglia from paraventricular area, cerebrocortex, and basal ganglion from the same batch of animals (left). In particular, the separation procedure for paraventricular tissue under microscopy is shown (right). (**B**) Volcano plot shows statistical significance (-log10) against log2 fold change when comparing the microglia derived from the above three anatomic area. (**C**) The common upregulated (left) and downregulated (right) genes when comparing the microglia derived from one area with the other two areas. The related gene numbers, as well their percentages among the total DEGs, are shown. (**D**) Top 10 KEGG annotations based on *P* values for the common DEGs of PvtMG when compared to the microglia from cerebrocortex and basal ganglion. The annotations associated with neurodegenerative diseases (purple asterisks) and proteolysis (red circles) are highlighted. (**E**) The average rankings regarding the expression abundance of *ApoE* among the 220 common upregulated DEGs between PvtMG and the other two areas according to gene expression values (TPM). (**F**) APOE expression, as indicated as volume ratio of positive signal over each cell, is compared between PvtMG and parenchymal microglia. Maximum z-projection of 37 μm. Arrowhead, PvtMG. Arrow, parenchymal microglia. Each dot represents the quantification from one 200 × 200 μm^2^ fields of view (FOVs), n = 6. \*\*\*\**P*<0.001 by two-tailed unpaired t test. Data are depicted as mean±SEM. For F, data are derived from 3 independent experiments.

All of the examined cells highly expressed marker genes of microglia, validating their identity (Figure S3B). Differential gene expression analysis identified hundreds of differentially expressed genes (DEGs) between microglia of different regions (Figure 4B).

However, there were more upregulated DEGs in the PvtMG when compared to the microglia of CTX or BG than those between CTX and BG microglia (Pvt vs. CTX: 410, Pvt vs. BG: 353, CTX vs. BG: 110, BG vs. CTX 177). In particular, PvtMG outnumbered their counterparts derived from the other two regions in common upregulated DEGs (220 vs. 51 in the CTX vs. 64 in the BG) (Figure 4C), suggesting that the PvtMG possess a region (ventricle)-specific transcriptomic signature. Kyoto Encyclopedia of Genes and Genomes (KEGG) analysis revealed that genes upregulated in PvtMG were primarily enriched for neurodegenerative diseases, including amyotrophic lateral sclerosis, Parkinson disease, Huntington disease, and Alzheimer disease (*Apoe*, *Psmb4*, *Kif5b*, *Rb1cc1*, *Psma7*, *Pik3r4*, *Adrm1* and *Cox6a1*) (purple asterisk in Figure 4D). In particular, *Apoe* ranked the top of the common upregulated DEGs in the PvtMG (Figure 4E). Our immunohistostaining verified the higher rate of APOE expression in the PvtMG than the nearby BG microglia (Figure 4F). In alignment with the upregulated pathways associated with neurodegenerative diseases, pathways associated with proteolysis were enhanced (red circle in Figure 4D), including those relevant to proteasome (*Asb13, Ube2d2a*, *Adrm1* and *Rnf187*), and autophagy (*Ykt6/Rps27a/Rb1cc1/Atg16l1/Pik3r4/Vps18*). Only 9 genes were commonly downregulated in the PvtMG relative to the microglia in the other examined regions, and all the suppressed pathways contained no more than one gene so they were not notable (Figure 4C).

### Acute loss of paraventricular microglia results in ventriculomegaly

It is tempting to speculate that the spatial positioning of PvtMG at ventricle walls endows the cells with particular roles. To interrogate the function of PvtMG, we designed a bead-based depletion strategy which only affect PvtMG, sparing parenchymal microglia (Figure 5A). The ependymal barrier precluded the permeabilization of 1 μm beads into the brain parenchyma. Employing this property, we infused fluorophore-labeled latex beads covalently coupled with diphtheria toxin (DT) into the lateral ventricle of *Cx3cr1*^CreERT2/+^:*iDTR* mice. Indeed, it showed that there was no beads permeabilized into the surrounding parenchyma (Figure 5B). Two days after DT-bead administration, almost all PvtMG at the lateral, 3^rd^, and 4^th^ ventricles were depleted, while parenchymal microglia were intact (Figure 5C). As expected, the number of CpMØ in stroma was normal since they were not exposed to the ventricles (Figure S4A). Very few beads appeared in the subarachnoid space (Figure S4B). In alignment, meningeal macrophages appeared unaffected in this period (Figure S4C).

**Figure 5.**
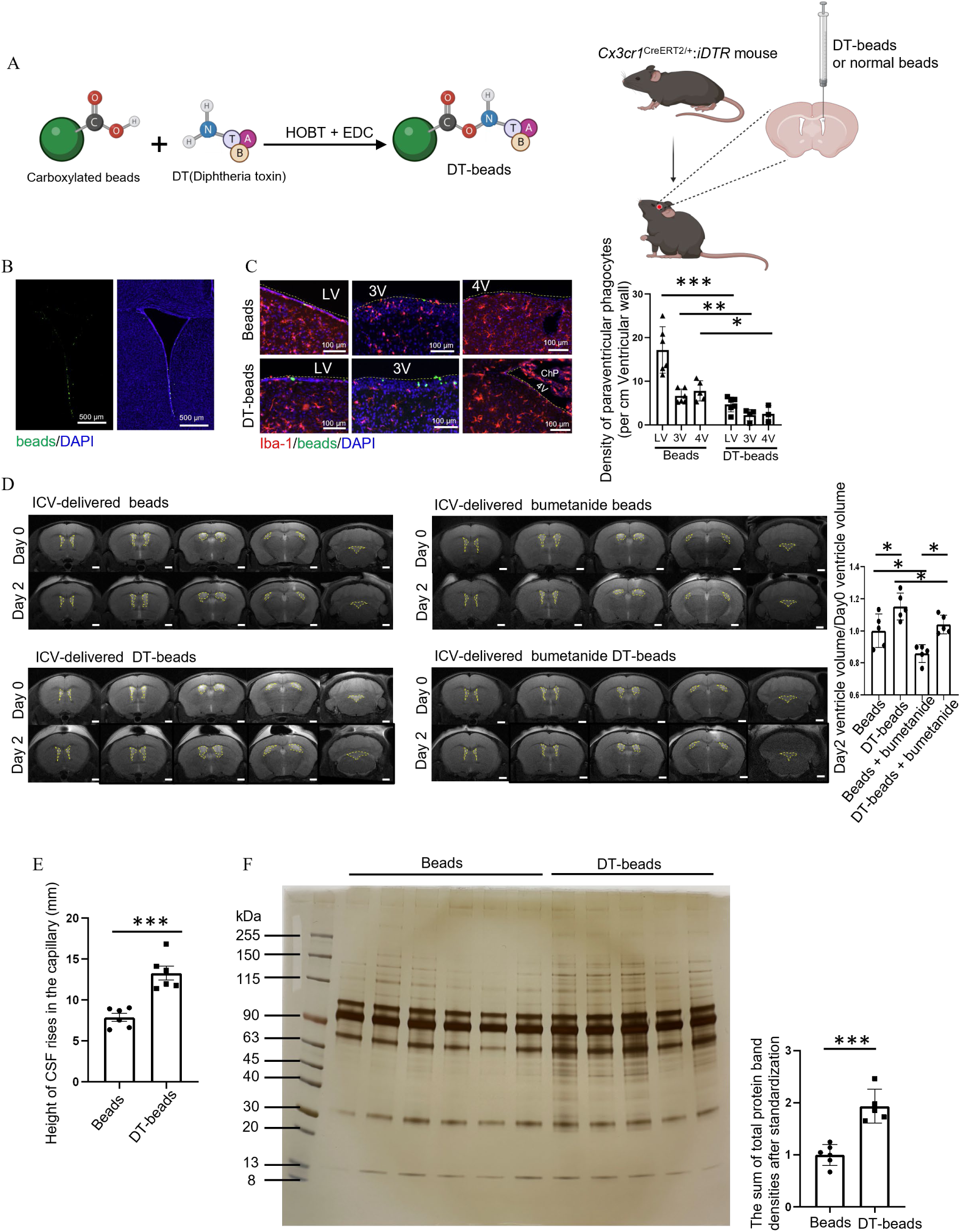
Depletion of PvtMG results in an increased colloid osmotic pressure and ventriculomegaly. (**A**) Schematic plot of ligation of DT and latex beads, as well as ICV injection of beads. (**B**) Representative images show the distribution of the ICV-injected fluorescent beads at 48 hr post injection. (**C**) Depletion of PvtMG in the wall of the indicated ventricles at 48 hr post DT-beads injection. (**D**) Representative MRI images of Representative MRI images of displayed sections are at bregma 0.6 mm, bregma -0.1 mm, bregma -0.6 mm, bregma -1.34 mm, and bregma -5.9 mm. The statistical comparison was conducted on the ventricular volume at the space within 1 mm anterior and posterior to the bregma -0.1 mm level. The ventricle areas were outlined by yellow dot lines. The ratio of the volume of two lateral ventricles on day 2 post bead injection versus the volume before bead injection of the same mice. During bead injection, some mice were also ICV infused with bumetanide. (**E**) Relative intraventricular hydraulic pressure, as measured by the rise of CSF in capillary after capillary was inserted into the cisterna magna for 5 s. (**F**) Silver staining for the proteins in the 2 μl CSF collected on Day 2 post the indicated ICV bead treatment. The total protein quantity was measured as the sum of band densities. \**P*<0.05, \*\**P*<0.01, \*\*\**P*<0.005 by two-tailed unpaired t test (C, E, F) and one-way ANOVA test (D). Data are depicted as mean±SEM.

Surprisingly, there was a noticeable expansion of the cerebral ventricles (ventriculomegaly) on day 2 post PvtMG depletion (Figure S5), as revealed by Hematoxylin and Eosin, which was confirmed by in vivo magnetic resonance imaging (MRI) (Figure 5D). Ventriculomegaly was accompanied by an elevated intraventricular hydraulic pressure (Figure 5E), suggesting an increased CSF volume. Acute ventriculomegaly could arise from impaired CSF efflux (e.g., due to obstruction of the cerebral aqueduct) and/or augmented choroid plexus-dependent CSF production. To examine CSF outflow in our PvtMG-depletion model, we injected the CSF tracer Evans Blue ^20^ into the lateral ventricles on day 2 and day 7 post DT-bead infusion and found that at both time points, dye distribution was unimpeded through the cerebral aqueduct and 4^th^ ventricle in 30 min (Figure S6), suggestive of aqueductal patency and preservation of bulk CSF flow. We next measured the effect of PvtMG depletion on the rate of CSF generation by ICV-delivered bumetanide, an inhibitor of choroid plexus-mediated CSF secretion ^20^. As expected, bumetanide administration reduced the ventricle volumes of both DT-bead-treated mice and bead-treated control mice; however, the ventricle volume of PvtMG-depleted mice were still larger than that of controls (Figure 5D). Thus, neither CSF blockade nor CSF over-production could explain the acute ventriculomegaly exhibited in the PvtMG-depleted mice. Next, we investigated the hypothesis that an increased CSF osmolality was responsible for the ventriculomegaly. To this end, we first examined the crystal osmotic pressure by measuring electrolytes and glucose in the CSF. It showed that there was no alteration of the concentrations of sodium, potassium, chloride or glucose in the CSF of PvtMG-depleted mice (Figure S7). We next examined the colloid osmotic pressure which was determined by protein concentrations. Quantitating total CSF protein by silver staining demonstrated a marked increase of protein in the CSF of PvtMG-depleted mice relative to the PvtMG-intact mice (Figure 5F). Together, the data above indicate an accumulation of possible “junk” protein in the CSF in the absence of PvtMG, which is in accordance with a strengthened proteolysis function of PvtMG, as suggested by the RNA-seq data.

## DISCUSSION

In this study, we identified paraventricular microglia, designated as PvtMG, as a subset of microglia distinguishable from parenchymal microglia. Populated in proximity to the ependymal wall of ventricles, PvtMG extended shorter processes in a lower density in comparison to their parenchymal counterparts. In particular, many processes derived from PvtMG were transependymal and were exposed to the apical CSF-contacting surface of the ventricles. Moreover, two-photon microscopy observation revealed that their processes had a movement polarity towards the ventricles, and even more pronouncedly, they would make transependymal migration to the apical side of ventricle walls. This observation strongly suggests that the previously recognized supraependymal phagocytes^21^ were derived the transmigrated PvtMG, which is consistent with the finding that in the microglia-deficient *Csf1r*^ΔFIRE/ΔFIRE^ mice, supraependymal and free-floating macrophages were completely lost while stromal CpMØ were only reduced^22^. The distinctive anatomical localization and unique activities well position the PvtMG as the sentinel cells monitoring the CSF in the ventricles. Indeed, we were able to show that these phagocytes were proficient in sequestering and phagocytosing ICV delivered exogenous particles, such as latex beads and red blood cells. It is feasible that the unique microenvironment of ventricle wall, i.e., the direct exposure to the CSF and/or a close contact with ependymal cells, may provide the differentiation cues for their identity, which requires further investigation.

The CSF is an integral part of the CNS and is essential for brain function and development. Thus, variation in its components can profoundly affect brain physiology. On the other hand, an abnormal brain function can affect the CSF composition, as seen in several neurological disorders^23,24^. In this perspective, regular drainage and reabsorption of CSF by the arachnoid villi and granulations is important, which, however, is not enough for a complete cleaning as demonstrated by our current study. We showed here that removing PvtMG would acutely have more protein accumulated in the CSF, suggesting that PvtMG critically regulate the biological environment of the CSF. Indeed, recent findings disclosed that a population of juxtatubular macrophages in the kidney^25^ and a lactation-induced macrophage subpopulation in the mammary gland^26^ committed to preventing tubular obstruction, suggesting almost all tubule systems in the body equipped with professional phagocytes as their plumbers for cleaning. Traditional AD biomarkers, such as β-amyloid 42 (Ab42), total Tau (Tau), and phosphorylated tau181 (pTau), are present in CSF and have demonstrated their analytical validity^27^. Future studies including proteomics should be set to disclose the composition alteration in the CSF in the absence of the PvtMG, which will shed light on not only what these cells are dedicated to cleaning, but regular “junks” generated by CNS metabolism as well.

Intraventricular hemorrhage (IVH) is a type of intracranial hemorrhage characterized by bleeding in the cerebral ventricular system, which would frequently lead to post-hemorrhagic hydrocephalus due to obstruction of intraventricular CSF flow^28^. Our observation that the PvtMG could efficiently phagocytose the ICV-injected red blood cells warrants future study of their roles in the pathogenesis and prognosis of IVH.

## Materials and Methods

### Mice

*Cx3cr1*^GFP/+^ mice (005582), *Cx3cr1*^CreERT2^ mice (020940, *Rosa26-*iDTR mice (007900), *Rosa26-stop-TdTomato* mice (007914) were from The Jackson Laboratory. *Ccr2*-deficient C57BL/6JSmoc-*Ccr2*^em1Smoc^ mice were purchased from Shanghai Model Organisms Center. All the mice were in C57BL/6 background. Normal C57BL/6 mice were purchased from Shanghai Research Center for Model Organisms. All mice used in this study without specific explanation were 8-to 12-week-old males. Mice were housed in a standard animal facility, with a 12-hr light/dark cycle, in specific-pathogen-free environment. All animal experiments adhered to the NIH Guide for the Care and Use of Laboratory Animals, and were approved by the Institutional Animal Care and Use Committee at Zhejiang University.

When treating the genetically inducible mice, tamoxifen dissolved in corn oil was i.p. injected into the mice with a daily dose of 75 μg/g body weight for 5 consecutive days.

### Intracerebroventricular (ICV) injection

Adult mice were anesthetized with isoflurane from a vaporizer. Following a midline scalp incision to expose the skull, the lateral ventricle was located by stereotaxic coordinates (A/P: 0.6 mm; M/L: 1.15 mm; D/V: 2.45 mm from the skull). A single delivery path was prepared above the lateral ventricle, followed by drilling a hole at the dimension of 0.5×0.5 mm using a micro-drill (Saeyang, Korea) to gain injection access into the left cerebral ventricle. Injection was aided by a nanoliter injector (Nanoject IIITM, Drummond Scientific, USA, Catalog#68018).

### Preparation of DT-conjugated latex beads and ICV injection

Two microliters of 1 μm carboxylated latex beads (Sigma-Aldrich; L4655) were diluted to 40 μl in ACSF (artificial cerebrospinal fluid)(Coolaber, SL6630-500ml), followed by co-incubation with 160 μl ACSF containing 100 μg of DT (List labs; 150) in a light-protected tube with continuous vortex overnight at room temperature. After that, beads were washed with cold PBS for three times to remove unbound free DT. Finally, the DT-conjugated beads were resuspended in 40 μl PBS. BSA-conjugated beads were used as controls. Beads were injected ICV at a volume of 5 μl per mouse at 1 μl min^−1^.

### Immunostaining and confocal imaging

Mice were anesthetized and transcardially perfused consecutively with cold PBS for 5 min and 4% paraformaldehyde (PFA) for another 5 min. Brain were then removed, fixed overnight in 4% PFA, and then dehydrated in PBS containing 30% sucrose at 4°C for 24 hr. Frozen coronal sections (30 µm thick) were made on a cryostat (SM2010R; Leica) at -20 °C. Sections were blocked for 2 hr at room temperature in a blocking buffer (5% donkey serum and 1% Triton X-100 in PBS). The sections were then incubated with primary antibodies in antibody dilution buffer (0.5% donkey serum and 0.1% Triton X-100 in PBS) overnight at 4 °C, followed by incubation with secondary donkey antibodies in antibody dilution buffer for 2 hr at room temperature in the dark. Primary antibodies used were anti-Iba1 (Wako; 019-19741, 011-27991), anti-CD206 (Bio-Rad; MCA2235T), anti-P2Y12R (Servicebio; GB112164), anti-TMEM119 (Abcam; ab209604), anti-β-catenin (Invitrogen; PA5-16762), anti-TER119 (Invitrogen; 14-5921-81), anti-apolipoprotein E (Abcam; ab183597), anti-Brdu (Biolegend; 364103), anti-AQP4 (Sigma-Aldrich; AB3594), anti-GFP (Abcam; Ab13970). After mounting on glass slides, stained sections were viewed under confocal microscope equipped with 10×/0.45, 20× /0.8, 40× /0.95, 63× /1.4 NA Plan-Apochromat objectives (Carl Zeiss LSM 900). Images (1024×1024 pixel resolution) were displayed as maximum-intensity projections of 25-28 µm thick Z stacks recorded by 1µm Z interval. Image analysis was performed using IMARIS software (Bitplane, Switzerland).

For H&E staining, brain samples were embedded in paraffin, sectioned into 5 µm thick slices, and subjected to H&E staining. The lateral ventricle area at the bregma 0.02 mm level was measured using OLYMPUS OlyVIA (version 3.1) software and adjusted to the area of the ipsilateral half of the brain.

### Preparation and two-photon imaging of live brain coronal sections

*Cx3cr1*^GFP/+^ mice were euthanized and the brains were harvested and kept on ice in a typical pre-oxygenated ACSF. The brains were sliced into 150 µm coronal sections with a Leica VT1000 S vibrating blade microtome (Leica Biosystems) at speed 1.80 and frequency 9. The brain sections were then stained with the aforementioned anti-AQP4 antibody (1:250) in ACSF at 37℃ for 50 min, followed by incubation with a fluorophore-conjugated secondary antibody for another 30 min to label paraventricular ependymal cells. Sections were held down with tissue anchors in 35 mm glass dishes and were imaged by Olympus FVMPE-RS two-photon microscope equipped with an environmental chamber and a motorized stage. Microscope configuration was set up for four-dimensional analysis (x, y, z, t). The fields were simultaneously recorded with an XLPlan N 25×/1.05 water immersion objective lens every 4 min for 2.5 hr with 1.5 µm Z axis increments and 800×800 pixel resolution. The imaging depth was ∼75 µm, with a Z-stack range of 25–30 µm. The laser was tuned to 920-nm excitation and used for all studies. Images and videos were processed using IMARIS (Bitplane) software.

### Isolation of periventricular microglia

Live brain sections obtained from *Cx3cr1*^GFP/+^ mice were quickly transferred to sterile slides, which were placed on a pre-cooled microscope stage. A 33-gauge needle was used to separate 30∼50 μm-wide ventricular walls under a 10 × objective. The separated tissues were immediately placed in pre-cooled ACSF and centrifuged at 300 g at 4°C. After the supernatant was discarded, 50 µl of DNase I (0.6 U/ml) diluted in ACSF was added to the tissues. The tissues were gently pipetted and mixed, then incubated at 37°C for 25 min. Dead cells were then labeled with 7-AAD. Live GFP^+^ microglia were purified by cell sorting with a BD SORP ARIA II.

### RNA-seq library construction

We analyzed 3 biological samples, each including 25 microglia from 2 mice. To this end, Holo-Seq, a recently developed method designed for improving data fidelity of low-cell-number samples^25,29,30^, to construct RNA-seq library. BBDuk (BBMap version 38.86) was utilized to remove adapters and trim low-quality bases from the raw sequencing reads. During this process, Illumina adapters and sequence regions with an average quality score below 15 were systematically eliminated. Subsequently, reads shorter than 36 base pairs were discarded. Following trimming, we aligned the left reads to the mouse genome (GRCm38, Genecode) using STAR (version 2.7.5a_2020-06-19) with the default parameters in paired-end mode. Finally, we quantified the transcript abundance by employing the ‘quant’ step of Salmon (version 1.2.1) and obtained 0.68±0.1 million genome mapped reads in basal ganglia samples, 1.34±0.05 million in paraventricular samples and 1.12±0.7 million in cerebrocortex samples, and within each group we detected 8429±365 genes, 9880±665 genes and 11117±343 genes respectively.

### RNA-seq data analysis

With the gene expression abundance data of microglia from three different brain regions, we utilized the R package DESeq2 (version 1.34.0)^31^ to conduct differential gene expression analysis between each pair of regions. During this process, we retained genes that had more than 10 mapped reads in at least two samples within either group, and identified significantly expressed genes with the |log2FC| > 1 and P value < 0.05 as cutoff. Subsequently, we performed GO analysis on these differentially expressed genes (DEGs) using the clusterProfiler package (version 4.0.5)^32^. The enrichGO function was employed with the following parameters: ‘OrgDb’ set to ‘org.Mm.eg.db’, ‘ont’ set to ‘BP’ (Biological Process), ‘pvalueCutoff’ set to 0.05, ‘qvalueCutoff’ set to 0.05, and ‘pAdjustMethod’ specified as ‘BH’ (Benjamin and Hochberg correction). To streamline the results, we utilized the simplify function to eliminate redundant GO terms, using a cutoff of 0.7, with ‘by’ set to “p.adjust”, and ‘select_fun’ set to min.

### Ventricle size acquisition by MRI (Magnetic Resonance Imaging)

#### MRI acquisition

Magnetic resonance imaging (MRI) was conducted on a Bruker 7.0T scanner (Bruker, PharmaScan 70/16, Germany), using a volume coil with an inner diameter of more than 23mm for transmission and a 4-channel phase-array coil for reception. Following a localizer scan for mouse positioning, an automatic global shimming was performed based on a pre-scanned B0 field map, and the shimming quality was confirmed by rechecking the field map. Anatomical images were acquired using a T2-weighted TurboRARE sequence in the axial plane with echo time (TE) = 35ms, repetition time (TR) = 2500ms, field of view (FOV) = 15×15mm^2^, matrix size = 256×256, slice thickness = 1mm, 15 slices, 2 averages, and the total scan duration was 2min 40s.

#### Data processing

For each mouse, the whole brain region was segmented using the Otsu thresholding method on a consistently positioned slice across different time points. K-means clustering (n=3) followed by morphological operations was applied to segment the ventricular region, while non-ventricular brain parenchymal areas were manually corrected using ITK-SNAP software (version 3.8.0, http://www.itksnap.org) by a scientist (JZ with 3 years of MRI experience) blinded to the task conditions to ensure unbiased data processing. Finally, the total number of pixels of the lateral ventricle was counted to assess changes in ventricle size.

### Examination of CSF generation and flow in the ventricular system

To detect whether the ventricular system was blocked, 5 μl Evans Blue (Sigma Aldrich; 0.5% in sterile ACSF) was ICV injected at a speed of 1 μl min^−1^. Five minutes after completion of the injection, the mice were sacrificed and the distribution of Evans Blue in each ventricle of the mice was examined. To examine the generation of CSF, bumetanide (Sigma-Aldrich, B3023) was dissolved in a Beads/DT-beads solution to a final concentration of 2.7 mM and administered at an injection rate of 1 µL/min, with a total injection volume of 5 µL per mouse. This was performed to observe the effect of choroid plexus cerebrospinal fluid production on ventricle size.

### CSF collection and analysis

Mice were anesthetized with isoflurane. The fur of the neck was shaved and cleaned with 70% iodine. Then, mice were placed in a stereotaxic frame to maintain the head fixed, and an ophthalmic solution was applied to prevent drying eyes. The skin from the neck was longitudinally incised and muscles were retracted using hooks to expose the cisterna magna. The glass capillary was pulled to a fine tip using a Micropipette Puller and mounted on a capillary holder attached to a micromanipulator. Under a dissecting microscope, the sharpened tip of the capillary was carefully broken with fine forceps to achieve an inner diameter of approximately 10–20 µm. Using the micromanipulator, the capillary tip was advanced near the dura mater at the cisterna magna until resistance was felt, ensuring that the membrane was not punctured. At this point, the capillary was tilted 30° toward the mouse tail from the vertical position. The micromanipulator controls were then used to gently guide the capillary through the dura mater, allowing CSF (10∼15 μl) to be automatically drawn into the capillary. This process would ensure very limited contamination from other tissues including the blood. The collected CSF was centrifuged at 1,500 g for 10 min at 4 °C to remove any cellular components and the supernatants were analyzed in this study.

### CSF pressure measurement

A capillary was inserted through the dura mater at the cisterna magna, as mentioned above. Once the CSF was observed in the capillary, we waited for 5 s before measuring the height of the CSF in the capillary. CSF heights in the collecting capillary were recorded and compared, which reflects the differences of CSF pressure between groups.

### SDS-PAGE electrophoresis of cerebrospinal fluid protein samples and silver stain

CSF sample were loaded onto SDS-PAGE gels. The proteins were electrophoresed at 170V for 40 min. Then, silver stain was conducted according to the manufacturer’s instructions (Beyotime; P0017S). ImageJ was used to calculate the density of bands.

### Statistics

Statistical analysis was performed with Prism 6.0 (GraphPad). Data are presented as mean ± SEM. One-way ANOVA with Tukey’s multiple-comparisons testing was used to compare multiple groups. Unpaired Student’s t tests were used to compare two groups. All statistical tests were two tailed, and P values of <0.05 were considered significant.

## Acknowledgments

We thank Qin Han for her technical assistance on Two-photon Microscopy from the Cryo-Electron Microscopy (CCEM) Center of Zhejiang University. We acknowledge the technical support provided by the Core Facility, Zhejiang University School of Medicine. We also thank Huang Yingying from the Core Facilities, Zhejiang University School of Medicine, for her technical support in primary microglia dissociation for bulk RNA-seq. This work was supported by grants from Zhejiang Province’s Vanguard Geese Leading Plan Project (2025C02151); Medical Health Science and Technology Project of Zhejiang Provincial Health Commission (WKJ-ZJ-2340); the Special Support Program for High Level Talents of Zhejiang Province (2022R52038); the National Natural Science Foundation of China (32170894 and 32470946 to X.Z.S, and 32471157 and 82170441 to P.S.).

## REFERENCES

1 Mack, A. F., Bihlmaier, R. & Deffner, F. Shifting from ependyma to choroid plexus epithelium and the changing expressions of aquaporin-1 and aquaporin-4. The Journal of physiology 602, 3097–3110, doi:10.1113/jp284196 (2024).

2 Theologou, M. et al. Cerebrospinal Fluid Homeostasis and Hydrodynamics: A Review of Facts and Theories. European neurology 85, 313–325, doi:10.1159/000523709 (2022).

3 Zhang, M., Hu, X. & Wang, L. A Review of Cerebrospinal Fluid Circulation and the Pathogenesis of Congenital Hydrocephalus. Neurochemical research 49, 1123–1136, doi:10.1007/s11064-024-04113-z (2024).

4 Oresković, D. & Klarica, M. The formation of cerebrospinal fluid: nearly a hundred years of interpretations and misinterpretations. Brain research reviews 64, 241–262, doi:10.1016/j.brainresrev.2010.04.006 (2010).

5 Süße, M., Saathoff, N., Hannich, M. & von Podewils, F. Cerebrospinal fluid changes following epileptic seizures unrelated to inflammation. European journal of neurology 26, 1006–1012, doi:10.1111/ene.13924 (2019).

6 Lehmann, S. et al. Cerebrospinal fluid A beta 1-40 peptides increase in Alzheimer’s disease and are highly correlated with phospho-tau in control individuals. Alzheimer’s research & therapy 12, 123, doi:10.1186/s13195-020-00696-1 (2020).

7 Barthélemy, N. R. et al. A soluble phosphorylated tau signature links tau, amyloid and the evolution of stages of dominantly inherited Alzheimer’s disease. Nature medicine 26, 398–407, doi:10.1038/s41591-020-0781-z (2020).

8 Da Mesquita, S. et al. Functional aspects of meningeal lymphatics in ageing and Alzheimer’s disease. Nature 560, 185–191, doi:10.1038/s41586-018-0368-8 (2018).

9 Harrison, I. F. et al. Impaired glymphatic function and clearance of tau in an Alzheimer’s disease model. Brain : a journal of neurology 143, 2576–2593, doi:10.1093/brain/awaa179 (2020).

10 Masuda, T. et al. Specification of CNS macrophage subsets occurs postnatally in defined niches. Nature 604, 740–748, doi:10.1038/s41586-022-04596-2 (2022).

11 Van Hove, H. et al. A single-cell atlas of mouse brain macrophages reveals unique transcriptional identities shaped by ontogeny and tissue environment. Nature neuroscience 22, 1021–1035, doi:10.1038/s41593-019-0393-4 (2019).

12 Goldmann, T. et al. Origin, fate and dynamics of macrophages at central nervous system interfaces. Nature immunology 17, 797–805, doi:10.1038/ni.3423 (2016).

13 Vidal-Itriago, A. et al. Microglia morphophysiological diversity and its implications for the CNS. Frontiers in immunology 13, 997786, doi:10.3389/fimmu.2022.997786 (2022).

14 Coates, P. W. Supraependymal cells: light and transmission electron microscopy extends scanning electron microscopic demonstration. Brain research 57, 502–507, doi:10.1016/0006-8993(73)90157-1 (1973).

15 Kenkhuis, B. et al. Co-expression patterns of microglia markers Iba1, TMEM119 and P2RY12 in Alzheimer’s disease. Neurobiology of disease 167, 105684, doi:10.1016/j.nbd.2022.105684 (2022).

16 Cui, J., Xu, H. & Lehtinen, M. K. Macrophages on the margin: choroid plexus immune responses. Trends in neurosciences 44, 864–875, doi:10.1016/j.tins.2021.07.002 (2021).

17 Drieu, A. et al. Parenchymal border macrophages regulate the flow dynamics of the cerebrospinal fluid. Nature 611, 585–593, doi:10.1038/s41586-022-05397-3 (2022).

18 Gerganova, G., Riddell, A. & Miller, A. A. CNS border-associated macrophages in the homeostatic and ischaemic brain. Pharmacology & therapeutics 240, 108220, doi:10.1016/j.pharmthera.2022.108220 (2022).

19 Farhadian, S. F. et al. Single-cell RNA sequencing reveals microglia-like cells in cerebrospinal fluid during virologically suppressed HIV. JCI insight 3, doi:10.1172/jci.insight.121718 (2018).

20 Robert, S. M. et al. The choroid plexus links innate immunity to CSF dysregulation in hydrocephalus. Cell 186, 764–785.e721, doi:10.1016/j.cell.2023.01.017 (2023).

21 Bleier, R., Albrecht, R. & Cruce, J. A. Supraependymal cells of hypothalamic third ventricle: identification as resident phagocytes of the brain. *Science (New York*, N.Y.) 189, 299–301, doi:10.1126/science.1145204 (1975).

22 Munro, D. A. D. et al. CNS macrophages differentially rely on an intronic Csf1r enhancer for their development. Development (Cambridge, England) 147, doi:10.1242/dev.194449 (2020).

23 Seino, Y. et al. Cerebrospinal Fluid and Plasma Biomarkers in Neurodegenerative Diseases. Journal of Alzheimer’s disease : JAD 68, 395–404, doi:10.3233/jad-181152 (2019).

24 Wang, S. Y. et al. Neurofilament Light Chain in Cerebrospinal Fluid and Blood as a Biomarker for Neurodegenerative Diseases: A Systematic Review and Meta-Analysis. Journal of Alzheimer’s disease : JAD 72, 1353–1361, doi:10.3233/jad-190615 (2019).

25 He, J. et al. Renal macrophages monitor and remove particles from urine to prevent tubule obstruction. Immunity 57, 106–123 e107, doi:10.1016/j.immuni.2023.12.003 (2024).

26 Cansever, D. et al. Lactation-associated macrophages exist in murine mammary tissue and human milk. Nature immunology 24, 1098–1109, doi:10.1038/s41590-023-01530-0 (2023).

27 Oboudiyat, C. et al. Cerebrospinal fluid markers detect Alzheimer’s disease in nonamnestic dementia. Alzheimer’s & dementia : the journal of the Alzheimer’s Association 13, 598–601, doi:10.1016/j.jalz.2017.01.006 (2017).

28 Symss, N. P. & Oi, S. Theories of cerebrospinal fluid dynamics and hydrocephalus: historical trend. Journal of neurosurgery. Pediatrics 11, 170–177, doi:10.3171/2012.3.peds0934 (2013).

29 Xiao, Z. et al. Holo-Seq: single-cell sequencing of holo-transcriptome. Genome Biol 19, 163, doi:10.1186/s13059-018-1553-7 (2018).

30 Wei, B. et al. Microglia in the hypothalamic paraventricular nucleus sense hemodynamic disturbance and promote sympathetic excitation in hypertension. Immunity 57, 2030–2042 e2038, doi:10.1016/j.immuni.2024.07.011 (2024).

31 Patro, R., Duggal, G., Love, M. I., Irizarry, R. A. & Kingsford, C. Salmon provides fast and bias-aware quantification of transcript expression. Nat Methods 14, 417–419, doi:10.1038/nmeth.4197 (2017).

32 Yu, G., Wang, L. G., Han, Y. & He, Q. Y. clusterProfiler: an R package for comparing biological themes among gene clusters. OMICS 16, 284–287, doi:10.1089/omi.2011.0118 (2012).

