## supplemental figures for "A distinctive subset of microglia positioned at the paraventricular zone are dedicated to cleaning the cerebrospinal fluid"

Figure S1

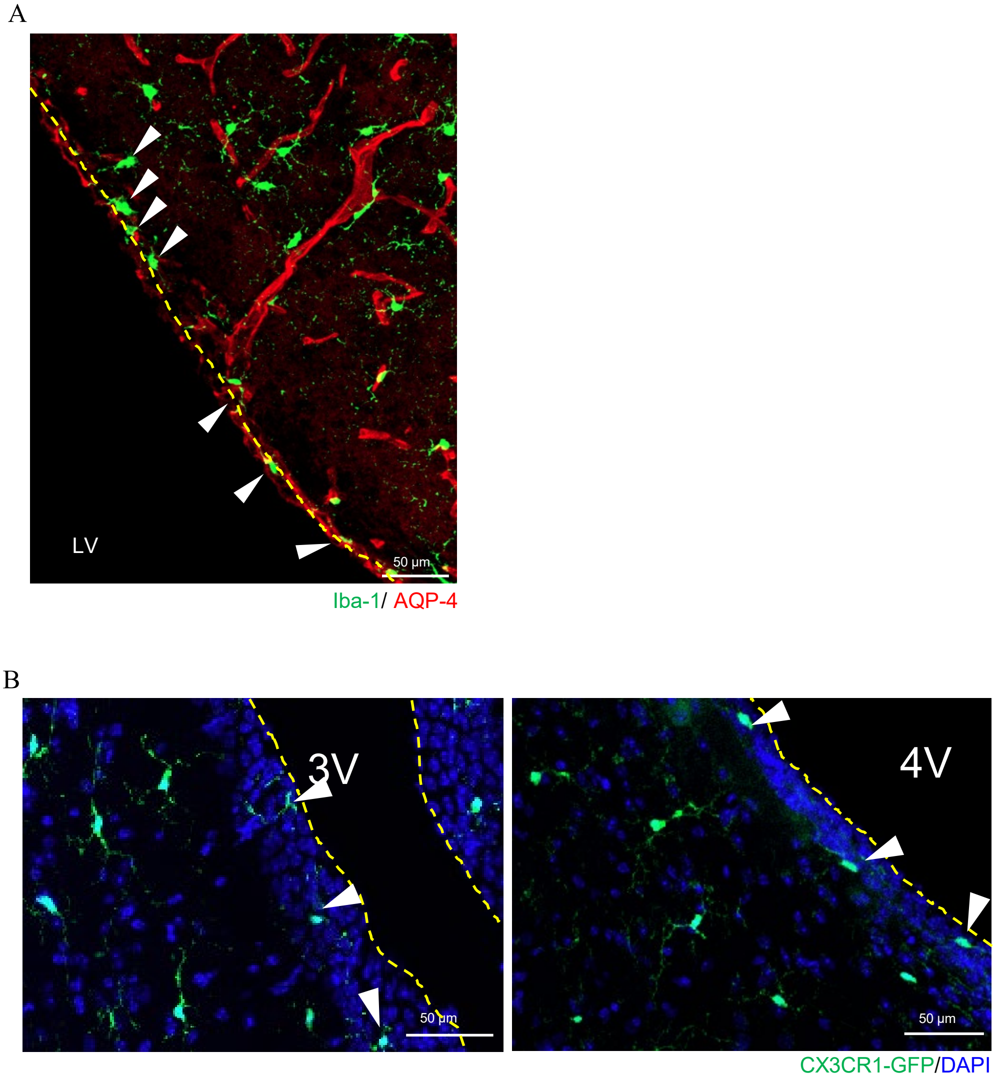


**Figure S1. A distinctive distribution pattern of paraventricular phagocytes.** (**A**) Immunohistochemical staining for Iba1 at the lateral ventricle (LV) wall area. (**B**) CX3CR1$^+$ cells are shown at the 3$^{rd}$ and 4$^{th}$ ventricle area indicated as 3V and 4V, respectively. The apical side of ventricular wall is indicated with dashed line.

Figure S2

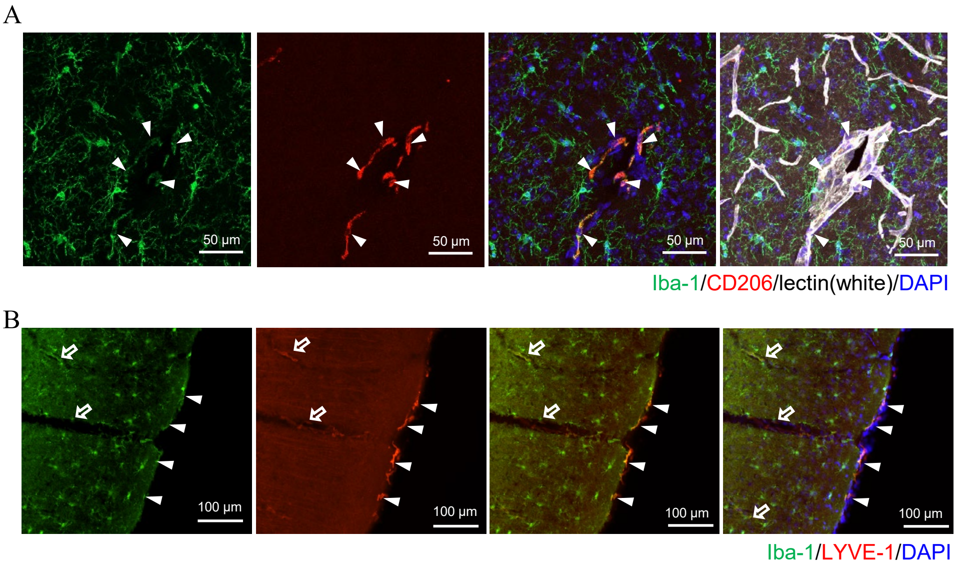


**Figure S2. Illustration of marker expression in brain border macrophages.** (**A**) Representative confocal images of cerebrocortical sections show that CD206 was expressed by perivascular macrophages ($Iba1^{+}$) at the wall of large non-capillary vessels. Vasculature was labeled by lectin. (**B**) Representative confocal images of LYVE-1 expression by $Iba1^{+}$ meningeal macrophages (arrowheads) and perivascular macrophages (arrows).

Figure S3

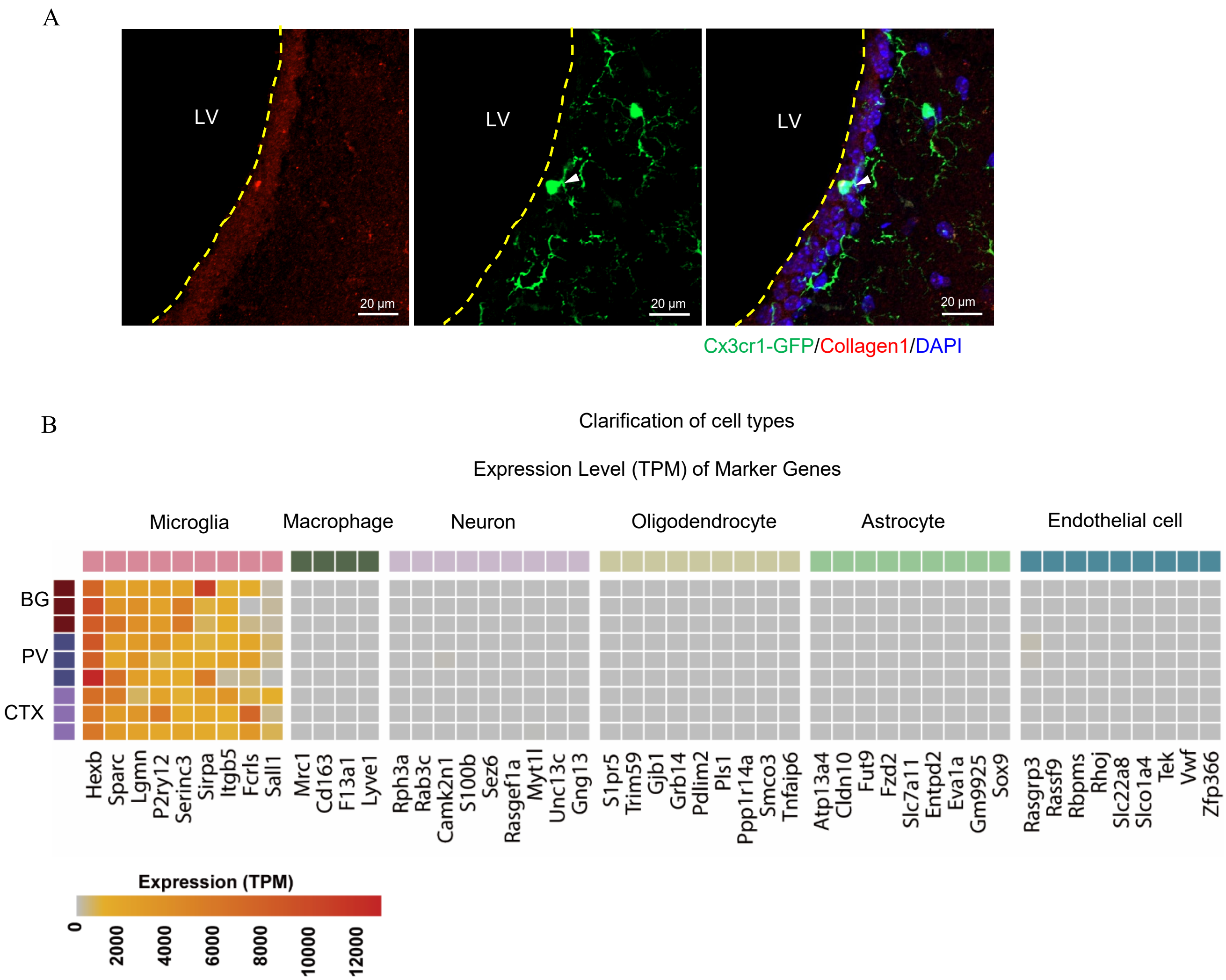


**Figure S3. Separating the paraventricular area and RNA-seq analysis.** (**A**) The ependymal wall of ventricles is enriched with collagen 1 deposition relative to the brain parenchyma. The apical side of ventricular wall is indicated with dashed line. (**B**) Based on normalized expression (TPM), heat map of the expression of characteristic genes of microglia, brain border macrophages, neurons, astrocytes, oligodendrocytes, and endothelial cells in the collected cells were shown according to the bulk RNA-seq data of microglia derived from basal ganglion (BG) and cerebrocortex (CTX) and the paraventricular phagocytes is shown.

Figure S4

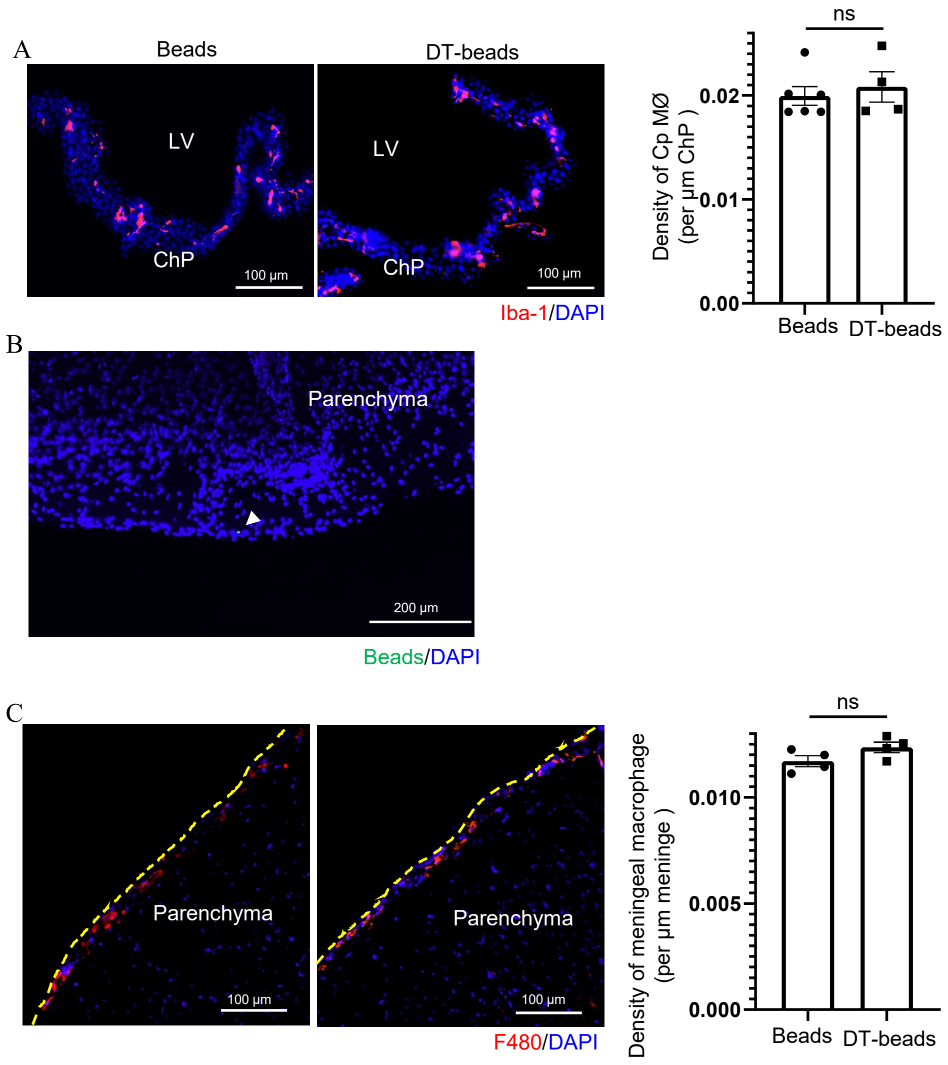


**Figure S4. CpMØ and meningeal macrophages are not affected in the *Cx3cr1*$^{CreERT2/+}$*:iDTR* mice by ICV administration of DT-beads.** (**A**) CpMØ had a normal density on day 2 post ICV DT-bead or normal bead treatment. ChP, Choroid plexuses. (**B**) Beads (green) were barely present in the subarachnoid space on day 2 post ICV DT-bead injection. (**C**) Meningeal macrophages had a normal density on day 2 post ICV DT-bead treatment. The meninge is indicated with dashed line. ns. not significant by two-tailed unpaired t test.

Figure S5

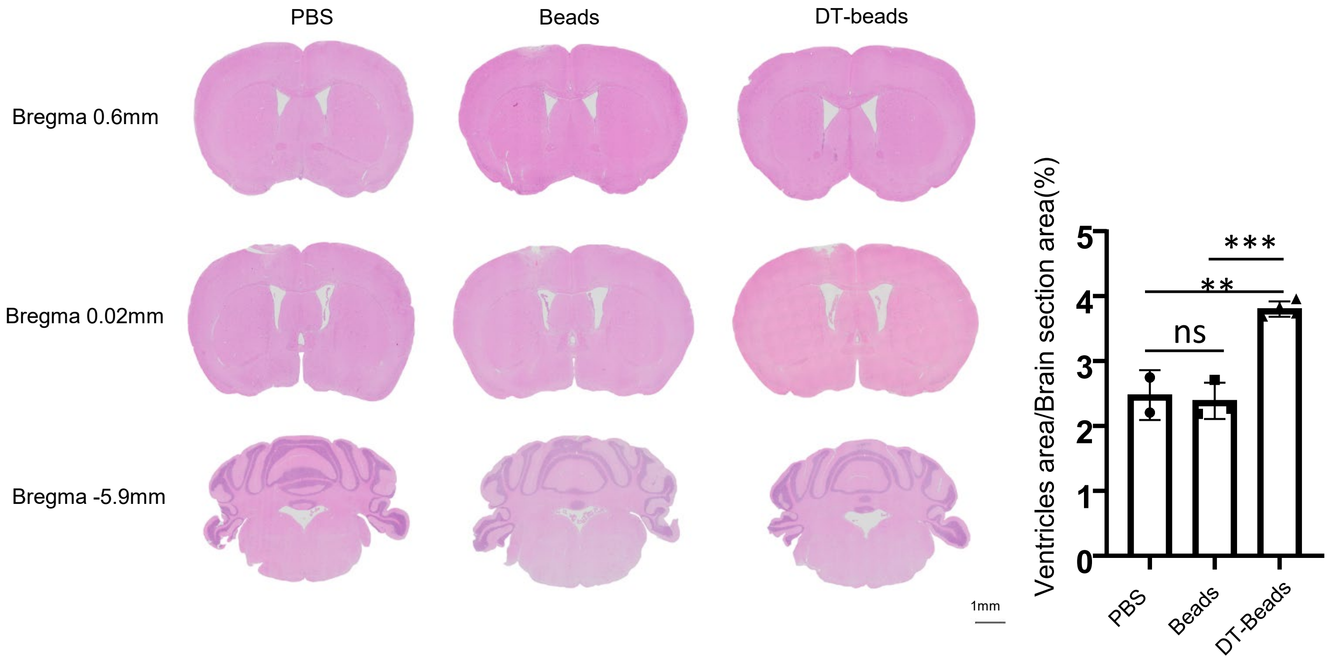


**Figure S5.** On day 2 post ICV treatment of PBS, normal beads or DT-beads, the brains of *Cx3cr1*$^{\mathrm{CreERT2/+}}$:*iDTR* mice were harvested and areas ranging from bregma -7.2 mm to 1.18 mm were subject to serial sectioning with a width of 5 μm. Representative H&E staining pictures show the section at bregma 0.6mm, bregma 0.02mm, and bregma -5.9mm of the same mice. The ventricle area at bregma 0.02mm was measured and compared between groups. ns, not significant. **P<0.01, ***P<0.005 by one-way ANOVA test. Data are depicted as mean±SEM. Data are derived from 2 independent experiments. The ventricle area at bregma 0.02mm was measured and compared between groups.

Figure S6

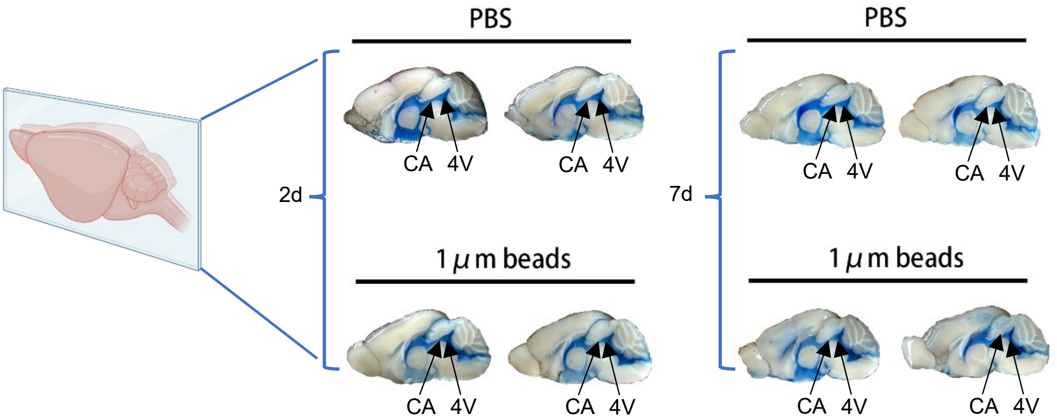


**Figure S6.** On day 2 and day 7 post ICV normal bead or DT-bead treatment, *Cx3cr1*$^{\mathrm{CreERT2/+}}$:*iDTR* mice received ICV infusion of 5 μl Evans Blue, and 30 min later, Evans Blue dispersion in the whole ventricular system including cerebral aqueduct (CA) and 4$^{\mathrm{th}}$ ventricle (4V) was examined.

Figure S7

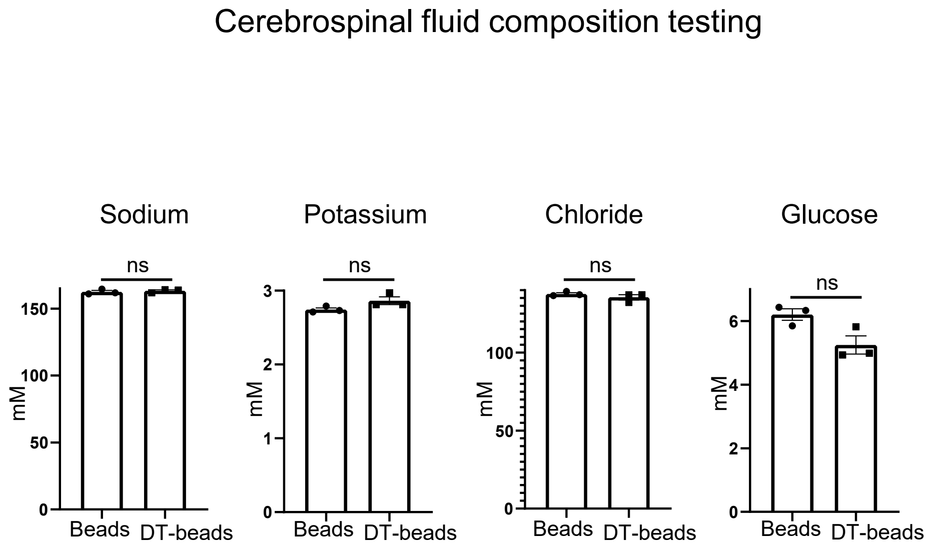


**Figure S7.** Two days post ICV injection of normal beads or DT-beads into *Cx3cr1*$^{\mathrm{CreERT2/+}}$:*iDTR* mice, the concentrations of sodium, potassium, chloride or glucose in the CSF were measured. ns, not significant by two-tailed unpaired t test.
