## Supplemental Video S1-S6 for "A distinctive subset of microglia positioned at the paraventricular zone are dedicated to cleaning the cerebrospinal fluid": 20250223 video legends.pptx

### Slide 1
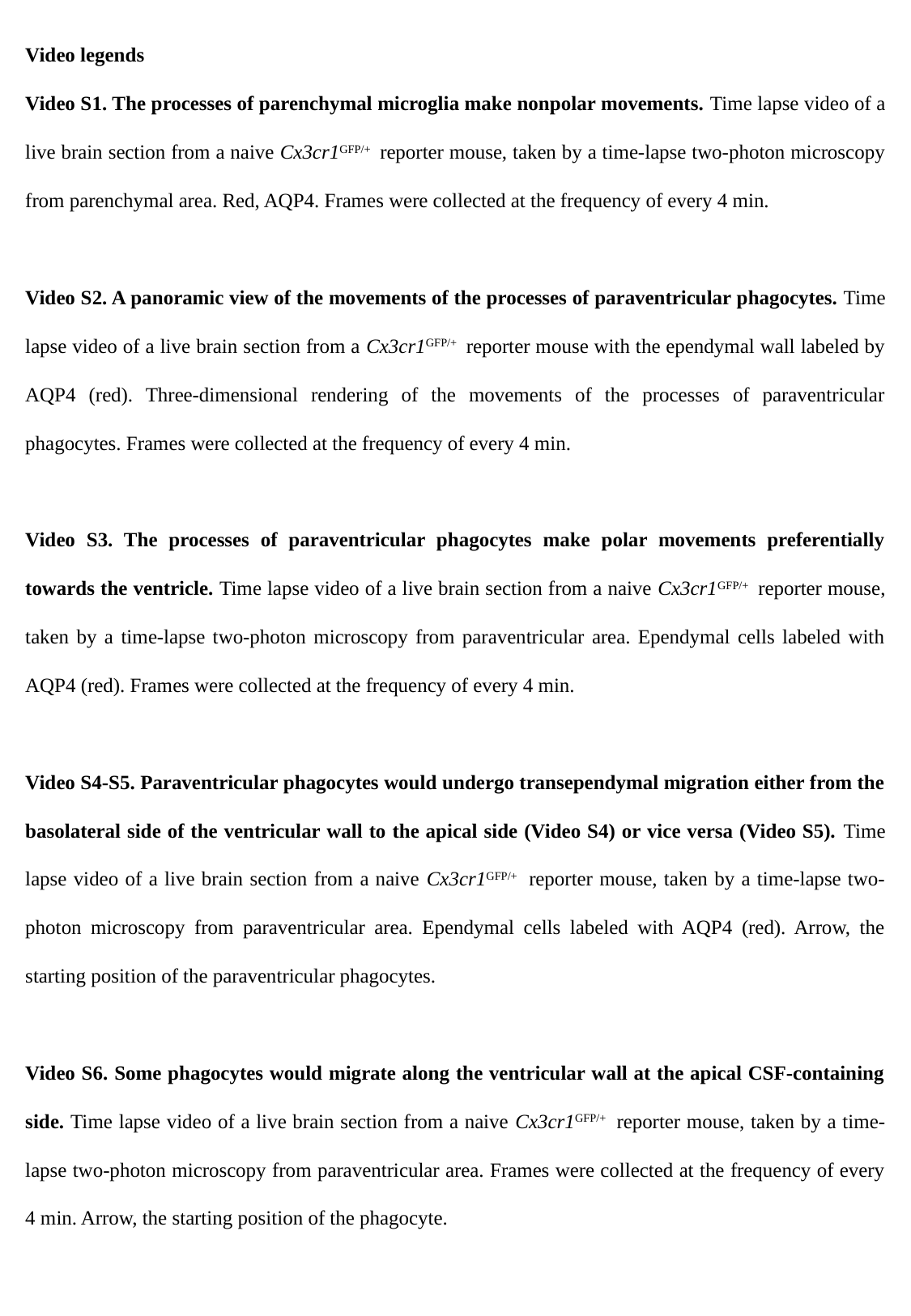

Video legends
Video S1. The processes of parenchymal microglia make nonpolar movements. Time lapse video of a live brain section from a naive Cx3cr1^GFP/+^ reporter mouse, taken by a time-lapse two-photon microscopy from parenchymal area. Red, AQP4. Frames were collected at the frequency of every 4 min.
Video S2. A panoramic view of the movements of the processes of paraventricular phagocytes. Time lapse video of a live brain section from a Cx3cr1^GFP/+^ reporter mouse with the ependymal wall labeled by AQP4 (red). Three-dimensional rendering of the movements of the processes of paraventricular phagocytes. Frames were collected at the frequency of every 4 min.
Video S3. The processes of paraventricular phagocytes make polar movements preferentially towards the ventricle. Time lapse video of a live brain section from a naive Cx3cr1^GFP/+^ reporter mouse, taken by a time-lapse two-photon microscopy from paraventricular area. Ependymal cells labeled with AQP4 (red). Frames were collected at the frequency of every 4 min.
Video S4-S5. Paraventricular phagocytes would undergo transependymal migration either from the basolateral side of the ventricular wall to the apical side (Video S4) or vice versa (Video S5). Time lapse video of a live brain section from a naive Cx3cr1^GFP/+^ reporter mouse, taken by a time-lapse two-photon microscopy from paraventricular area. Ependymal cells labeled with AQP4 (red). Arrow, the starting position of the paraventricular phagocytes.
Video S6. Some phagocytes would migrate along the ventricular wall at the apical CSF-containing side. Time lapse video of a live brain section from a naive Cx3cr1^GFP/+^ reporter mouse, taken by a time-lapse two-photon microscopy from paraventricular area. Frames were collected at the frequency of every 4 min. Arrow, the starting position of the phagocyte.
